# Genomic characterization of serial-passaged Ebola virus in a boa constrictor cell line

**DOI:** 10.1101/091603

**Authors:** Greg Fedewa, Sheli R. Radoshitzky, Xiǎolì Chī, Lián Dǒngb, Melissa Spear, Nicolas Strauli, Mark D. Stenglein, Ryan D. Hernandez, Peter B. Jahrling, Jens H. Kuhn, Joseph DeRisi

## Abstract

Ebola virus disease (EVD) is a viral hemorrhagic fever with a high case-fatality rate in humans. EVD is caused by four members of the filoviral genus *Ebolavirus*, with Ebola virus (EBOV) being the most notorious one. Although bats are discussed as potential ebolavirus reservoirs, limited data actually support this hypothesis. Glycoprotein 2 (GP2) of reptarenaviruses, known to infect only boa constrictors and pythons, are similar in sequence and structure to ebolaviral glycoprotein 2 (GP2), suggesting that EBOV may be able to infect snake cells. We therefore serially passaged EBOV and a distantly related filovirus, Marburg virus (MARV), in the boa constrictor kidney cell line, JK, and characterized viral growth and mutational frequency by sequencing. We observed that EBOV efficiently infected and replicated in JK cells, but MARV did not. In contrast to most cell lines, EBOV infected JK cells did not result in obvious cytopathic effect (CPE). Genomic characterization of serial-passaged EBOV in JK cells revealed that genomic adaptation was not required for infection. Deep sequencing coverage (>10,000x) demonstrated the existence of only a single non-synonymous variant (EBOV glycoprotein precursor preGP T544I) of unknown significance within the viral population that exhibited a shift in frequency of at least 10% over six passages. Our data suggest that boid snake derived cells are competent for filovirus infection without appreciable genomic adaptation; that cellular filovirus infection without CPE may be more common than currently appreciated; and that there may be significant differences between the natural host spectra of ebolaviruses and marburgviruses.

**IMPORTANCE:** Ebola virus (EBOV) causes a high case-fatality form of viral hemorrhagic fever. The natural reservoir of EBOV remains unknown. EBOV is distantly related to Marburg virus (MARV), which has been found in bats in the wild. The glycoprotein of a reptarenavirus known to infect boid snakes (pythons and boas) shows similarity in sequence and structure to these viruses, suggesting that EBOV and MARV may be able to infect and replicate in snake cells. We demonstrate that JK, a boa constrictor cell line, does not support MARV infection, but does support EBOV infection without causing overt cytopathic effect or the need for appreciable adaptation. These findings suggest different filoviruses may have a more diverse natural host spectra than previously thought.

## INTRODUCTION

Ebola virus (EBOV) is one of five members of the genus *Ebolavirus* in the mononegaviral family *Filoviridae*. Four ebolaviruses (Bundibugyo virus, EBOV, Sudan virus, Taï Forest virus) are known to cause Ebola virus disease (EVD), whereas the fifth member, Reston virus (RESTV), is thought to be nonpathogenic for humans. EVD is clinically indistinguishable from Marburg virus disease (MVD), which is caused by the two members of the filoviral genus *Marburgvirus* (Marburg virus [MARV] and Ravn virus [RAVV]) (1). The latest EVD outbreak, caused by EBOV, began in Western Africa in December 2013 and ended in March 2016, infecting 28,646 and killing 11,323 people (2). Like the vast majority of EVD outbreaks (2, 3), this outbreak started with a single introduction of EBOV from an unknown wild reservoir host into a human, with subsequent human-to-human transmission (4-10).

Frugivorous bats are often discussed as potential ebolaviral host reservoirs, but supporting data are overall sparse and stem largely from detection of anti-EBOV or anti-RESTV antibodies or short, EBOV genome-like, RNA fragments by RT-PCR. Ebolaviruses have not been recovered from any wild bat; ebolavirus genomes have not been sequenced from wild bats; and experimental infections of frugivorous bats with ebolaviruses have thus far failed (11-14). In contrast, genetically diverse MARV and RAVV could repeatedly be isolated from wild Ugandan Egyptian rousettes (*Rousettus aegyptiacus*), a frugivorous bat species, in direct vicinity of human infections (15, 16) and experimental infections of Egyptian rousettes have been successful in the laboratory (11). Together, these findings indicate that ebolaviruses and marburgviruses may differ in host tropism and that in contrast to marburgviruses, bats may not play a major role in ebolavirus maintenance in nature. However, until now, filovirus genus-specific cell susceptibility differences have not been uncovered *in vitro*, i.e., cells lines that can be infected with marburgviruses typically also support ebolavirus infection independent of species origin (3).

The recent discovery of a possible distant evolutionary relationship (17) between the glycoprotein genes of filoviruses and snake-infecting reptarenaviruses (*Arenaviridae*: *Reptarenavirus*) (16) prompted us to test the filovirus susceptibility of the boid snake (python and boas) cell line, boa constrictor JK (18) to evaluate whether filoviruses have the general ability to replicate in non-mammalian cells. We demonstrate that JK cells can be infected over multiple passages with EBOV, but not MARV; that EBOV infection of JK cells is not accompanied by cytopathic effect (CPE); and that EBOV does not undergo major genomic adaptation while replicating in this cell line. Our data support the hypothesis that there may be fundamental differences in ebolavirus and marburgvirus host tropism in the wild and indicate a need for further investigation of filovirus host tropism using non-mammalian cell lines.

## MATERIALS AND METHODS

### Filovirus stock preparation

Infections with Ebola Virus/H.sapiens-tc/COD/1995/Kikwit-9510621 (reference genome GenBank #KT582109; EBOV) (19) and Marburg virus/H.sapiens-tc/KEN/1980/Mt. Elgon-Musoke (MARV) (20) were conducted under biosafety level 4 conditions at the United States Army Medical Research Institute of Infectious Diseases (USAMRIID). EBOV and MARV were propagated in grivet (*Chlorocebus aethiops*) kidney epithelial Vero E6 cells (American Type Culture Collection, Manassas, VA, #CCL-81) and titrated by plaque assay as previously described (21-23).

### Quantification of filoviral titers by qRT-PCR

Boa constrictor kidney JK cells were plated at 15,000 per well in a 96-well plate as previously described (18). One day later, media were removed, and cells were infected with EBOV or MARV (MOI = 1 or 10) or mock infected (no virus) (50 μl/well). Inocula were removed 1 h later, and cells were washed once with phosphate-buffered saline (PBS) and supplemented with fresh growth media (150 μl/well). Cells were incubated at 37°C in a 5% CO2 atmosphere. At the indicated time points (0, 24, 48, 72, and 144 h after virus inoculation), media were either harvested for qRT-PCR or titer was determined by plaque assay (data not shown). At the experiment endpoint (144 h), cells were fixed with formalin (Val Tech Diagnostics, Pittsburgh, PA USA) for immunostaining. For qRT-PCR, RNA was extracted with Trizol (Thermo Fischer Scientific, Waltham, MA USA) and the Ambion Blood RNA Isolation Kit (Thermo Fischer Scientific, Waltham, MA USA). The assay was performed with RNA UltraSense one-step kit (Thermo Fisher Scientific Waltham, MA USA) and TaqMan probe (ABI, Thermo Fischer Scientific, Waltham, MA USA) following the manufacturer’s instructions. The primers used were: EBOGP_For (TGGGCTGAAAACTGCTACAATC), EBOGP_Rev (CTTTGTGCACATACCGGCAC), probe EBOGP_Prb (5-6FAM-CTACCAGCAGCGCCAGACGG-TAMRA) (24), and MARV_GP2_F (TCACTGAAGGGAACATAGCAGCTAT), MARV_GP2_R (TTGCCGCGAGAAAATCATTT), and probe MARV_GP2_P (ATTGTCAATAAGACAGTGCAC). Serial 10-fold dilutions (102 to 107) of the assayed virus were used as standards.

### Filovirus virus serial passage

EBOV or MARV were passaged in either JK cells or human epithelial adenocarcinoma HeLa cells (American Type Culture Collection #CCL2). For each of the serial passages, JK cells and HeLa cells were plated in six-well plates (at 300,000 cells/well, three replicates per cell line per virus). One day later, cells were exposed to EBOV or MARV at a multiplicity of infection (MOI) of 1. Briefly, exposure was performed by first removing media from cells, incubating cells with media containing filovirus for 1 h, washing cells, and finally adding fresh media back to cells. Infected cells were then incubated at 37°C in a 5% CO2 atmosphere for 3 or 4 days (Fig. 1). Supernatants were collected at the indicated time points; 50 μl were used to infect monolayers of fresh cells; and 1.5 ml were added to Trizol for sequencing.

**FIG 1:**
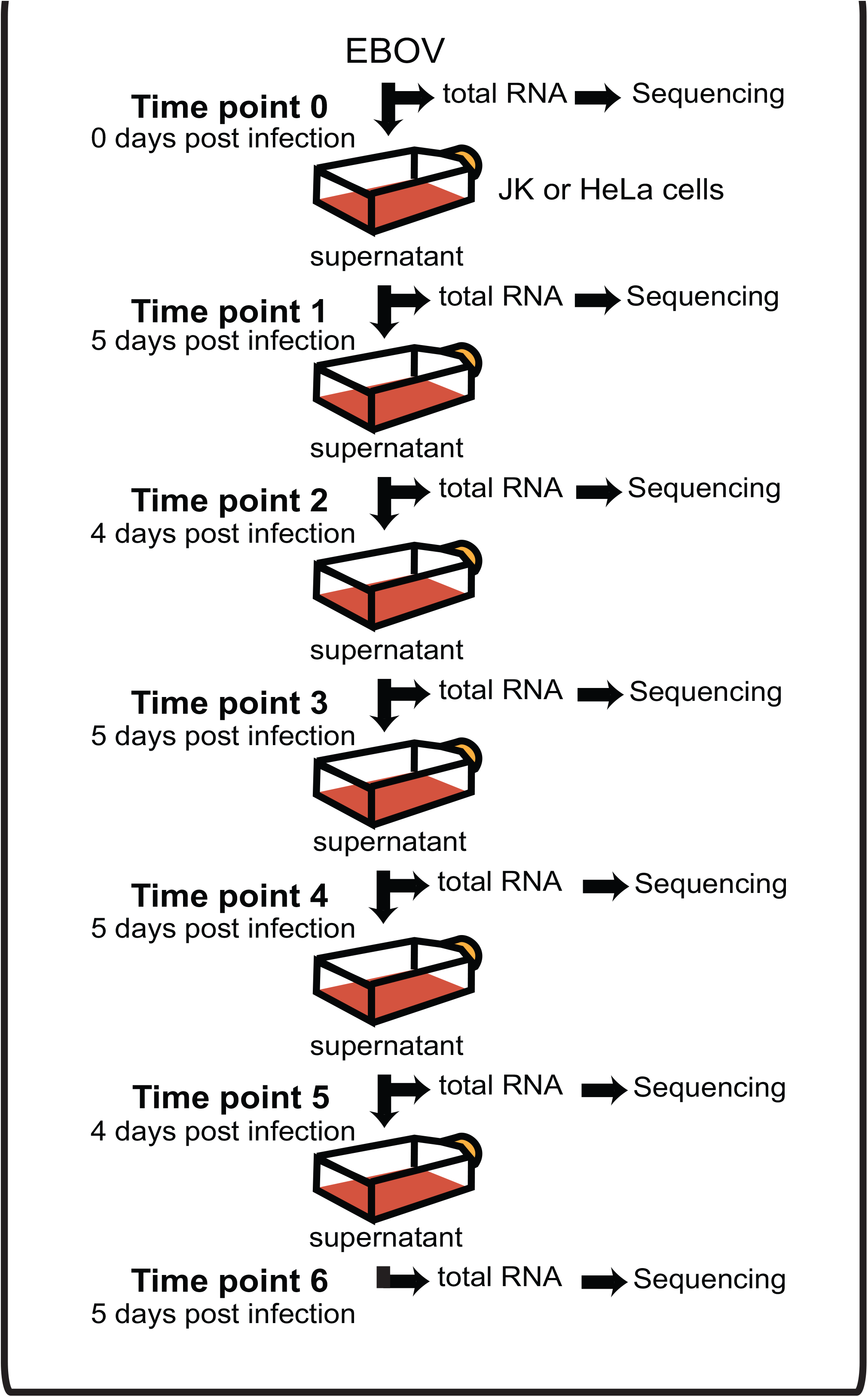
Cartoon of the viral passaging experimental procedure. Plated cells, either boa constrictor JK or human HeLa cells, were infected with EBOV for 1 h and then grown for either 4 or 5 days. To passage virus, supernatants were removed and a 1/40 subsample (50 μl) was used to inoculate a fresh monolayer of cells. In addition, 1.5 ml of the supernatant was inactivated for sequencing. This procedure was repeated for a total of 6 passages of EBOV.

### Filovirus immunostaining

Cells infected with EBOV or MARV were stained for high-content quantitative image-based analysis with murine monoclonal antibodies against EBOV or MARV GP1,2 (6D8 and 9G antibody, respectively), followed by Alexa Fluor 488-conjugated goat anti-mouse IgG (Invitrogen, Thermo Fisher Scientific, Waltham, MA USA). Infected cells were also stained with Hoechst 33342 and HCS CellMask Red (Invitrogen, Thermo Fisher Scientific, Waltham, MA USA) for nuclei and cytoplasm detection, respectively. Infection rates and cell numbers were determined using high-content quantitative imaging data on an Opera quadruple excitation high sensitivity confocal reader (model 3842 and 5025; PerkinElmer, Waltham, MA USA) at two exposures using ×10 air, ×20 water, or ×40 water objective lenses as described in (24). Analysis of the images was accomplished within the Opera environment using standard Acapella scripts. At least 4,000 cells and up to 9,000 cells, were analyzed per well.

### Measurement of Cytopathic effects

We measured cell number as an indication of CPE. See Filovirus immunostaining methods for details. Briefly, infected cells were also stained with Hoechst 33342 and HCS CellMask Red (Invitrogen, Thermo Fisher Scientific, Waltham, MA USA) for nuclei and cytoplasm detection, respectively. Infection rates and cell numbers were determined using high-content quantitative imaging data on an Opera quadruple excitation high sensitivity confocal reader.

### Passage population size measurement

The number of EBOV genomes that each passage produced and the number of genomes added to sequencing libraries were determined by two-step reverse transcription droplet digital PCR (RT-ddPCR) (25). EBOV RNA was reverse-transcribed using EBOV-specific primer EBOGP_For (TGGGCTGAAAACTGCTACAATC), diluted, and assayed with the Bio-Rad Qx200 Droplet Digital PCR System (Bio-Rad, Hercules, CA USA) following the manufacturer’s instructions.

### Sequencing-library preparations

Trizol inactivated samples were prepared for Illumina sequencing using a protocol slightly-modified from our previously published protocol (26) Briefly, complementary DNA (cDNA) was created from randomly primed RNA using SuperScript VILO Master Mix (Thermo Fisher Scientific, Waltham, MA USA). cDNA was tagmented using Illumina’s Nextera reagents (Illumina, San Diego, CA, USA), followed by dual-barcoding to prevent miscalling of samples (27). Libraries were quantified by qPCR, pooled, size-selected using BluePippin (Sage Science, Beverly, MA, USA), amplified, quantified again by qPCR, and paired-end sequenced (150/bases) on an Illumina HiSeq 4000 system at the University of California, San Francisco Center for Advanced Technology. Samples HeLa-P1-R1 (Host-Passage-Replicate) and JK-P1-R through JK-P6-R1 were prepared and sequenced separately using the same method and sequencer.

### Single nucleotide variant analysis pipeline

Sequencing reads were filtered for reads containing sequencing adapters and quality using a cut-off of at least 95% of the sequence having a 0.98 probability being correct (-rqf 95 0.98) with PriceSeqFilter from PRICE (version 1.2) (28). Filtered reads were aligned to the EBOV reference genome [KT582109 bases 1–18882] using GSNAP (version 2015-09-29) (29) using default settings.

Because of the very high coverage in each sample, duplicate reads were not removed, a step usually taken in single nucleotide variant (SNV) analysis. Sorted and indexed BAM files were processed with LoFreq* (version 2.1.2) (30) using default settings, to call SNVs. A final cut-off of ≥0.005 allele frequency was selected as a conservative threshold, calculated as 1. standard deviations above the mean of each nucleotide’s maximum detected allele frequency (0.00339, σ = 0.00129) of the Illumina supplied PhiX control sequence, which was included in each sequencing run. SNVs were then determined to be either synonymous or non-synonymous. Analysis was performed and graphs were generated using Python3, IPython (31), pandas (32), matplotlib (33), and seaborn (34).

### Testing for selection

See supplemental methods. Briefly, we developed a simulation-based procedure to identify alleles in the EBOV genome that changed frequency over passages more than expected under neutrality given the dynamic viral population size and estimated sequencing error rates. The neutral simulations had five parameters: the overall population growth function, the number of generations, the starting allele frequency, and the read depth for each site during the first and last passage.

### Detection of defective interfering genomes

Sequencing reads were processed in the same way as for SNV analysis. For each passage point, only properly paired reads were used. All of the passages of replicate 1 in JK cells (JK-R1) and passage 1, replicate 1, of passage in HeLa cells (HeLa-R1-P1) had a sizable drop in Q-score during sequencing of read 2. These reads were filtered out during pre-processing, necessitating that these paired-end reads be mapped as a combined single-end sample for each of the above passages. These combined samples then lacked proper pairing and were not used in defective interfering (DI) genome analysis. Each of the properly paired reads were also confirmed for the correct mapping orientation. Then the “reference location” located in each samples’ BAM file was used as that read’s mapping location and the distance difference between the read 1 mapping location and read 2 mapping location was calculated along with the mean and standard deviation for the entire set. Proper pairs characterized by a distance difference greater than the mean + 3 σ were counted as reads coming from potential DI genomes.

### Data availability

Sequencing data from EBOV passaging is located on NBCI SRA under BioProject: PRJNA353512.

## RESULTS

### Ebola virus, but not Marburg virus, replicates in boa constrictor cells

To test whether filoviruses can replicate in boa constrictor cellsx, we exposed a previously established boa constrictor kidney cell line, JK (18), to either EBOV or MARV at MOIs of 1 or 10. At various time points after exposure, cell culture supernatant was collected for qRT-PCR, or cells were fixed and stained for filoviral antigen (GP1,2) detection (24). Based on immunostaining, 26.22% (σ = 4.56) and 60.98% (σ = 6.46) of the cells were infected with EBOV at 144 h post inoculation (hpi) at MOIs of 1 and 10, respectively. Mock-exposed cells were not infected (0.11%, σ = 0.08) as expected. Surprisingly, exposure to MARV resembled mock. Based on immunostaining, we measured MARV infection for the mock infection at 3.01% (σ=0.50), for an MOI of 1 at 7.20% (σ=2.22), and for an MOI of 10 at 12.57% (σ=2.69). Quantification of filoviral RNA by qRT-PCR corroborated the immunostaining assay results. For EBOV-infected JK cells, we measured 1.87 × 108 (σ = 2.30 × 107) and 8.46 × 108 (σ = 3.45 × 108) genome copies/ml at 144 hpi at MOIs of 1 and 10, respectively. This represents a 49-fold and 23-fold increase, respectively, over 0 h post inoculation. For MARV-infected JK cells, we measured 3.68 × 106 (σ = 3.41 × 106) and 2.43 × 107 (σ = 7.63 × 106) genome copies/ml at 144 hpi representing a 1.30-fold and 1.27-fold increase at MOIs of 1 and 10, respectively.

### Ebola virus does not cause cytopathic effects in JK cells

Cells were stained with Hoechst 33342, imaged and counted as an indication of cell viability. When compared to mock infected, EBOV-infected JK cells do not show a decrease in the number of viable cells, unlike what has been shown for many other cell lines (35). We counted 83,678 (σ = 292) cells per well of mock infected JK cells, while EBOV-infected cells were counted at 83,678 (σ = 546) cells per well for MOI of 1, and 6,539 (σ = 827) cells per well for MOI of 10.

### Ebola virus, but not Marburg virus, continues to replicate in boa constrictor cells during serial passage

To characterize any adaptive genomic mutations necessary for efficient growth in JK cells, we serially passaged EBOV in JK cells in parallel with control human (HeLa) cells for 6 cycles (an average of 4.33 days per cycle) (Fig. 1) and MARV, analogously, for 5 cycles. The infection of both JK and HeLa cells was initiated at an MOI of 1 (3.0 x10^5^ plaque forming units (pfu)/well). For each passage cycle of EBOV, the extent of infection was monitored by qRT-PCR, immunostaining, and reverse transcription digital-droplet PCR (RT-ddPCR). For each passage cycle of MARV, the extent of infection was monitored by qRT-PCR. While EBOV was detected by qRT-PCR in both JK and HeLa cells at all passages, MARV was detected at all passages in HeLa cells, but only at the first passage in JK cells (Table S2). At all passages, EBOV infected JK cells revealed clusters of EBOV GP_1,2_-positive cells, with predominantly cytoplasmic and cell membrane staining (Fig. 2). Over the course of these passages, the number of genome equivalents produced by infected JK cells was modestly lower than by infected HeLa cells. Quantification of EBOV genome copy number in the supernatants from passages in JK cells by RT-ddPCR yielded an average of 8.49 x10^8^ copies/ml (σ = 9.92 x10^8^) across all passages and replicates, whereas HeLa cells yielded an average genome copy number of 6.34 x10^9^ copies/ml (σ = 5.88 x10^9^). The EBOV genome copy number measured in the JK supernatants was not significantly different between the first and last passage (4.34 x10^9^ vs. 1.79 x10^9^, p=0.4, Welch’s t-test).

**FIG 2:**
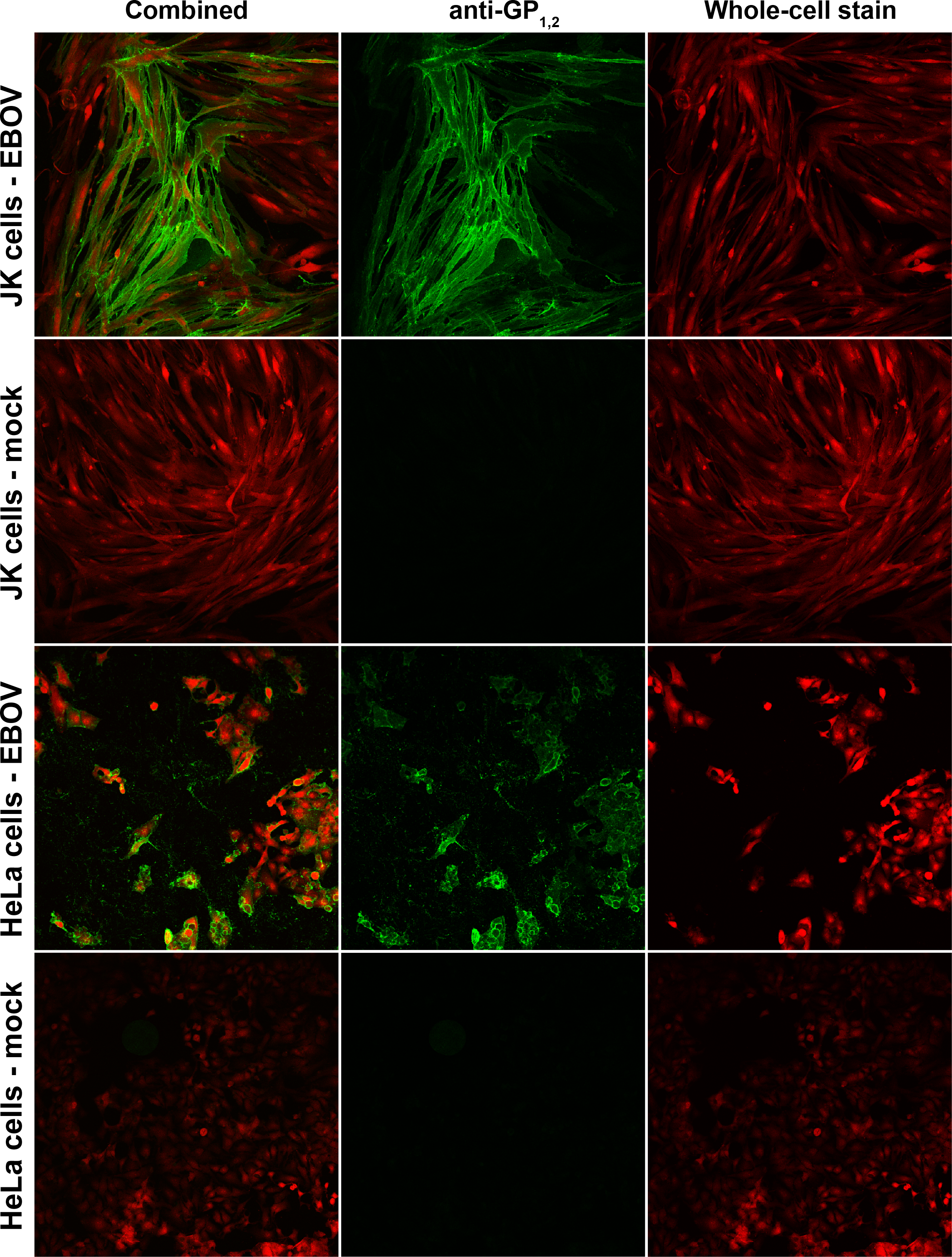
Antibody staining of EBOV GP1,2. Cells infected with EBOV were stained for cytoplasm (shown in red) and with anti-GP1,2 antibody (shown in green).

We used a deep sequencing approach to characterize the spectrum of possible mutations associated with EBOV adaptation to JK cells. For each passage, total cell culture supernatant RNA was processed into cDNA libraries for deep sequencing by random priming. For each library, sequencing reads were aligned to the EBOV reference genome. The mean coverage of the EBOV genome in JK cells across all passages was 36,730-fold (σ = 12,016), and 69,946-fold (σ = 26,582) for HeLa cell passages (Fig. 3). We detected no regional bias of coverage at any point within the genome in any of the three biological replicates for infected JK and HeLa cells, excluding the extreme 5’ and 3’ ends. Previous characterization of cells infected with either EBOV or MARV using deep sequencing yielded a pronounced gradient of filovirus gene transcription similar to that seen for other mononegaviruses. Transcripts accumulate in the 3’ to 5’ direction, with the furthest 3’ gene (encoding the filoviral nucleoprotein [NP]) yielding the highest coverage and the furthest 5’ gene (encoding the filoviral RNA-dependent RNA polymerase [L]) yielding the lowest coverage (36). For the data presented here, the lack of a 3’ to 5’ coverage gradient is consistent with sequence reads derived from EBOV genomic RNA in cell culture supernatant virions, as opposed to cellular EBOV transcripts (Fig. 3).

**FIG 3:**
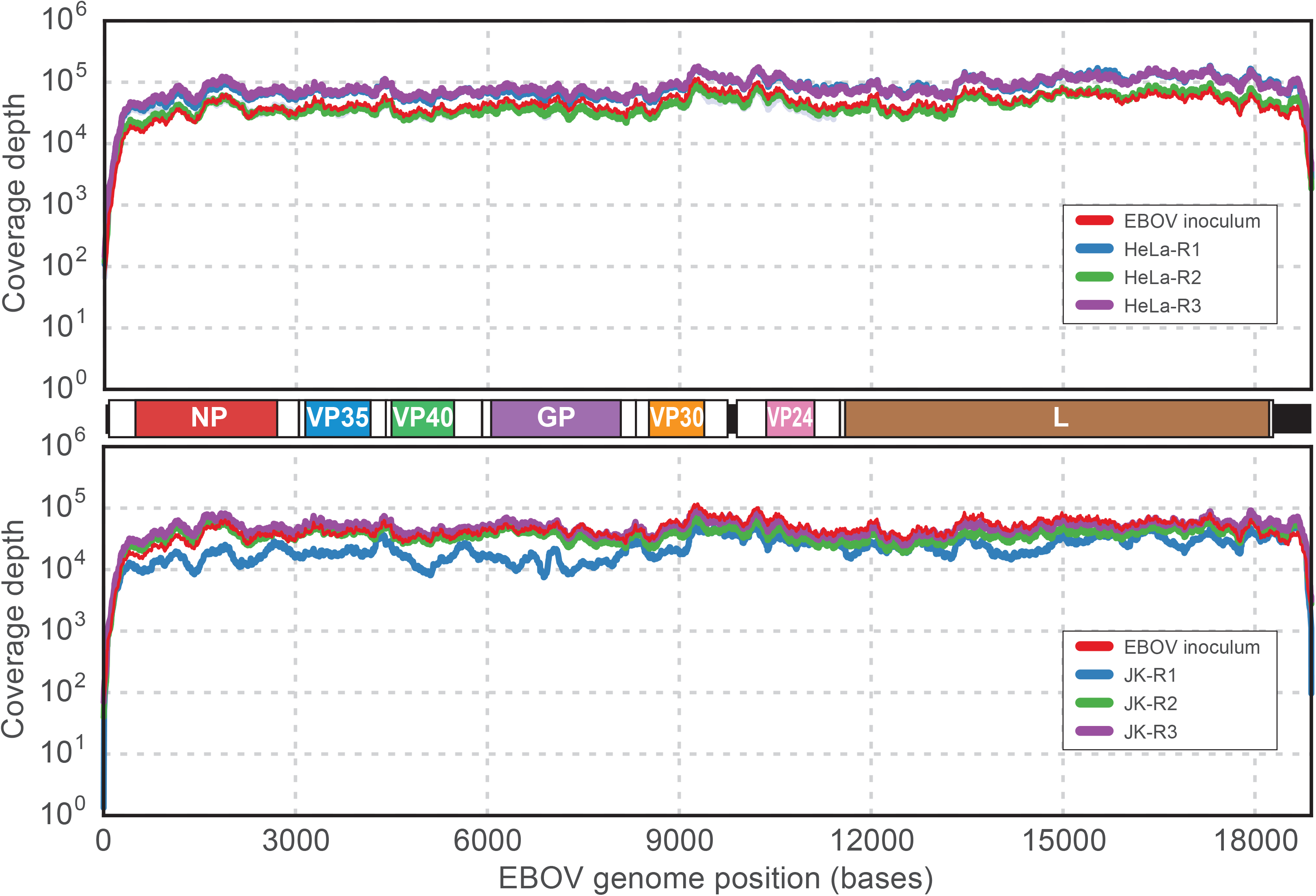
Coverage maps of mapped reads. Each sample was deep-sequenced and mapped back to the EBOV reference genome (genome cartoon drawn to scale between both maps). The number of reads that map to each genome base position was computed for each sample. For each replicate passage series for either HeLa (top graph: EBOV inoculum, red; HeLa-R1, blue; HeLa-R2, green; HeLa-R3, purple) or JK cells (bottom graph: EBOV inoculum, red; JK-R1, blue; JK-R2, green; JK-R3, purple), the mean coverage (respectively-colored solid lines) was calculated and graphed.

These data identify boa constrictor JK cells as susceptible to EBOV, but not MARV, infection. To our knowledge, JK cells represent the first cell line with filovirus genus-specific (ebolavirus vs. marburgvirus) susceptibility to infection.

### Ebola virus adaption is not required for infection of boa constrictor cells

We first characterized the extent of variation within the EBOV inoculum population. We detected 48 single nucleotide variants (SNVs) in the inoculum that passed our quality and frequency cut-off filters including 21 non-synonymous SNVs. We detected only a single position (nt 7669, EBOV glycoprotein precursor [preGP] codon 544: T544I) with a nonsynonymous SNV having an allele frequency of >10% in the inoculum (Table 1, Table 2, Fig. 4A). At this position, the initial population consisted of 62.0% (Thr) and 37.9% (Ile), similar to the previously characterized EBOV/Kik-951061 “R4414” (passage 2) strain (19).

**FIG 4:**
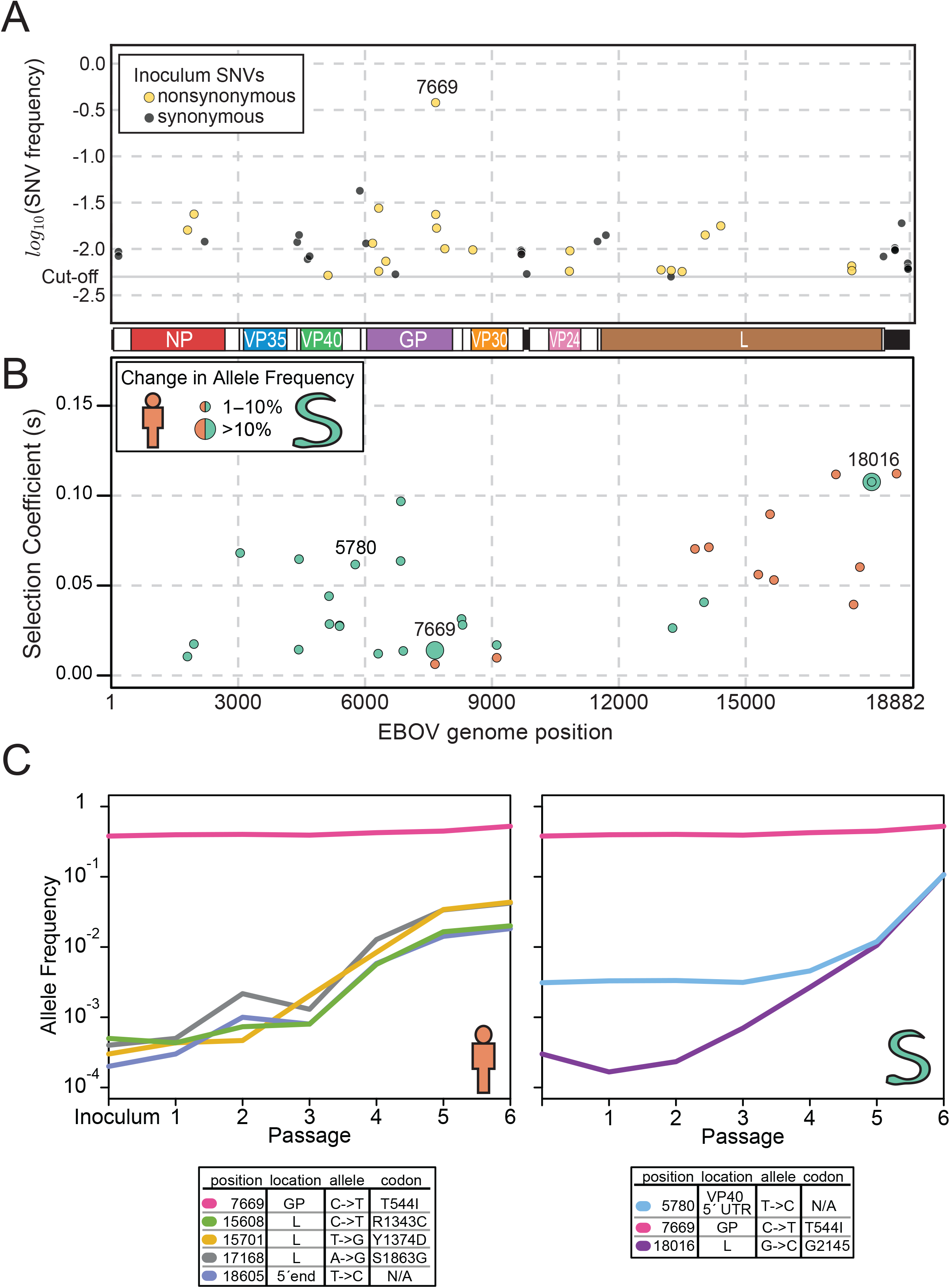
Alleles across the EBOV genome A: Single nucleotide variants (SNVs) found in the EBOV passaging inoculum. The log10 (allele frequency) of each SNV is plotted as a function of its position in the EBOV reference genome (genome cartoon drawn to scale between A and B). All SNVs are color coded. Yellow: non-synonymous SNVs; black: synonymous and non-coding SNVs. B: The estimated selection coefficients across the EBOV genome for passages in HeLa cells (orange) and JK cells (green). Each point represents the most positively selected allele for each site in the EBOV genome. Selection coefficients were averaged across the three replicates. C: The allele frequency trajectories across passages of the most strongly selected sites in HeLa (right) and JK (left) cells.

**Table 1:** EBOV passage data. Each sample is described as cell type used for EBOV passaging (Host); passage number (Passage); replicate number (Replicate); mean coverage (Mean coverage); total number of single nucleotide variants (SNVs) found for that sample (Total SNVs); number of non-synonymous SNVs found (Non-syn SNVs); number of coding-synonymous SNVs (Coding syn SNVs); number of SNVs found in each viral passage-replicate not found in the inoculum or below the limit of detection in the inoculum (Non-synSNVs not in inoculum); number of SNVs found in all three replicates, but not, or below the limit of detection, in the inoculum (Non-syn SNVs in all replicates number); number of EBOV genomes produced in each sample found by RT-ddPCR (Genome copies by RT-ddPCR); number of EBOV genomes produced per ml for each sample found by RT-ddPCR (Genome copies per ml by RT-ddPCR); number of EBOV genomes produced per ml for each sample found by RT-qPCR (Genome copies per ml by RT-qPCR); and the fraction of reads that have a putative deletion between reads (DI read fraction). N/A, not applicable.

|  |  |  | Mean coverage | Total SNVs | Non-syn SNVs | Coding Syn SNVs | Non-synSNVs not in inoculum | Non-syn SNVs in all replicates | Genome copies by RT-ddPCR | Genome copies per ml by RT-ddPCR | Genome copies per ml by RT-qPCR |  |
| --- | --- | --- | --- | --- | --- | --- | --- | --- | --- | --- | --- | --- |
| Host | Passage | Replicate |  |  |  |  |  |  |  |  |  | DI read fraction |
| Vero E6 | 0 |  | 46599 | 48 | 21 | 6 | N/A | N/A | 2.46E+08 | 4.92E+08 | N/A | 0.000276 |
| HeLa | 1 | 1 | 114414 | 55 | 26 | 5 | 0 | 0 | N/A | N/A | 1.06E+10 | N/A |
|  |  | 2 | 50683 | 113 | 52 | 19 | 14 |  | 3.11E+10 | 1.55E+10 | 1.14E+10 | 0.000286 |
|  |  | 3 | 114461 | 102 | 46 | 19 | 13 |  | 3.11E+10 | 1.55E+10 | 1.14E+10 | 0.000344 |
|  | 2 | 1 | 57860 | 70 | 32 | 9 | 5 | 1 | 3.84E+10 | 1.92E+10 | 6.60E+09 | 0.001306 |
|  |  | 2 | 42273 | 79 | 38 | 12 | 8 |  | 1.69E+10 | 8.47E+09 | 4.26E+09 | 0.001207 |
|  |  | 3 | 110186 | 86 | 36 | 12 | 7 |  | 2.86E+10 | 1.43E+10 | 5.39E+09 | 0.001629 |
|  | 3 | 1 | 84746 | 87 | 37 | 12 | 8 | 1 | 2.43E+08 | 1.22E+08 | 8.61E+09 | 0.001437 |
|  |  | 2 | 33734 | 54 | 21 | 8 | 2 |  | 1.27E+10 | 6.33E+09 | 3.49E+10 | 0.001587 |
|  |  | 3 | 90833 | 89 | 41 | 10 | 10 |  | 5.39E+09 | 2.70E+09 | 2.23E+10 | 0.001721 |
|  | 4 | 1 | 109716 | 80 | 35 | 13 | 10 | 5 | 1.97E+08 | 9.86E+07 | 1.34E+10 | 0.000351 |
|  |  | 2 | 51857 | 80 | 38 | 11 | 9 |  | 8.60E+09 | 4.30E+09 | 8.53E+09 | 0.000364 |
|  |  | 3 | 91247 | 80 | 35 | 10 | 7 |  | 9.15E+09 | 4.57E+09 | 7.36E+09 | 0.000314 |
|  | 5 | 1 | 79159 | 120 | 60 | 20 | 23 | 5 | 7.43E+09 | 3.72E+09 | 5.12E+09 | 0.000320 |
|  |  | 2 | 32086 | 147 | 76 | 37 | 51 |  | 5.59E+09 | 2.79E+09 | 5.61E+09 | 0.000368 |
|  |  | 3 | 65817 | 90 | 43 | 12 | 14 |  | 9.33E+09 | 4.66E+09 | 3.12E+09 | 0.000365 |
|  | 6 | 1 | 56325 | 42 | 25 | 7 | 12 | 3 | 1.91E+09 | 9.56E+08 | 9.17E+09 | 0.000512 |
|  |  | 2 | 49650 | 165 | 91 | 43 | 61 |  | 7.14E+09 | 3.57E+09 | 7.73E+09 | 0.000549 |
|  |  | 3 | 47332 | 57 | 30 | 7 | 14 |  | 1.89E+09 | 9.43E+08 | 6.98E+09 | 0.000595 |
| JK | 1 | 1 | 34972 | 71 | 27 | 8 | 4 | 0 | 1.71E+09 | 8.53E+08 | 2.59E+09 | N/A |
|  |  | 2 | 70411 | 40 | 21 | 4 | 2 |  | 9.31E+09 | 4.65E+09 | 2.72E+09 | 0.000210 |
|  |  | 3 | 103515 | 32 | 15 | 3 | 0 |  | 2.01E+09 | 1.01E+09 | 1.98E+09 | 0.000157 |
|  | 2 | 1 | 7237 | 39 | 20 | 6 | 6 | 0 | 7.08E+08 | 3.54E+08 | 7.00E+08 | N/A |
|  |  | 2 | 24005 | 25 | 13 | 1 | 0 |  | 1.25E+09 | 6.23E+08 | 3.88E+08 | 0.000208 |
|  |  | 3 | 24138 | 31 | 13 | 3 | 1 |  | 4.72E+08 | 2.36E+08 | 3.15E+08 | 0.000209 |
|  | 3 | 1 | 13131 | 33 | 15 | 3 | 2 | 1 | 5.03E+08 | 2.52E+08 | 6.97E+08 | N/A |
|  |  | 2 | 19078 | 28 | 16 | 3 | 1 |  | 7.68E+08 | 3.84E+08 | 8.26E+08 | 0.000289 |
|  |  | 3 | 20975 | 48 | 22 | 2 | 4 |  | 1.15E+09 | 5.76E+08 | 2.98E+10 | 0.000255 |
|  | 4 | 1 | 18038 | 58 | 25 | 9 | 8 | 1 | 1.39E+09 | 6.95E+08 | 6.70E+08 | N/A |
|  |  | 2 | 39866 | 49 | 22 | 8 | 8 |  | 2.50E+09 | 1.25E+09 | 6.78E+08 | 0.000215 |
|  |  | 3 | 66186 | 37 | 19 | 3 | 6 |  | 1.63E+09 | 8.15E+08 | 6.10E+08 | 0.000225 |
|  | 5 | 1 | 8475 | 71 | 32 | 18 | 12 | 0 | 3.07E+08 | 1.54E+08 | 2.28E+08 | N/A |
|  |  | 2 | 14266 | 66 | 33 | 13 | 18 |  | 1.22E+09 | 6.10E+08 | 2.58E+08 | 0.000326 |
|  |  | 3 | 16464 | 59 | 29 | 10 | 16 |  | 2.46E+08 | 1.23E+08 | 2.31E+08 | 0.000307 |
|  | 6 | 1 | 56183 | 98 | 41 | 24 | 23 | 2 | 1.33E+09 | 6.66E+08 | 1.31E+09 | N/A |
|  |  | 2 | 54147 | 64 | 33 | 9 | 18 |  | 9.81E+08 | 4.91E+08 | 6.21E+08 | 0.000188 |
|  |  | 3 | 60200 | 63 | 31 | 15 | 20 |  | 3.07E+09 | 1.54E+09 | 1.17E+09 | 0.000225 |
|  |  | Mean | 53521 | 69 | 33 | 11 | 15/8 (HeLa/JK) |  | 1.27E+10/1.70E+09 (HeLa/JK) |  |  | 0.00078/0.00023 (HeLa/JK) |

DI, Defective interfering; EBOV, Ebola virus; non-syn, nonsynonymous; SNV, single nucleotide variants; syn, synonymous;

**Table 2:** EBOV inoculum population sequence variation. Each of the SNVs we found above our cut-off in the inoculum population. Each row is the position (Position), the allele from the reference sequence used (Reference allele), the variant allele (SNV allele), the percent saturation of the variant allele (SNV %), the gene where the allele is located (Gene), the codon where the allele is located and the change it caused (Codon change), and the sequencing depth at that position (Sequencing Depth).

| Table 2: EBOV inoculum population sequence variation |  |  |  |  |  |  |
| --- | --- | --- | --- | --- | --- | --- |
| Position | Reference allele | SNV allele | SNV % | Gene | Codon change | Sequencing Depth |
| 170 | C | A | 0.93 | NP |  | 3219 |
| 172 | T | C | 0.84 | NP |  | 3217 |
| 1805 | C | T | 1.60 | NP | P446S | 51212 |
| 1958 | C | T | 2.38 | NP | P497S | 60425 |
| 2209 | T | C | 1.20 | NP | S580 | 29331 |
| 4397 | A | G | 1.18 | VP35 |  | 61244 |
| 4441 | C | T | 1.42 | VP40 |  | 52069 |
| 4643 | C | T | 0.78 | VP40 | A55 | 30954 |
| 4691 | A | G | 0.83 | VP40 | S71 | 28166 |
| 5125 | T | C | 0.52 | VP40 | I216T | 28914 |
| 5878 | T | G | 4.25 | VP40 |  | 36349 |
| 6023 | G | T | 1.15 | GP |  | 46354 |
| 6179 | G | T | 1.15 | GP | E47D | 47583 |
| 6324 | G | A | 2.75 | GP | V96M | 49964 |
| 6325 | T | C | 0.57 | GP | V96A | 46789 |
| 6493 | C | T | 0.74 | GP | A152V | 40212 |
| 6719 | C | A | 0.53 | GP | T227 | 49001 |
| 7669 | C | T | 37.95 | GP | T544I | 36890 |
| 7672 | A | C | 2.36 | GP | E545A | 35520 |
| 7692 | G | A | 1.68 | GP | D552N | 35084 |
| 7888 | A | C | 1.01 | GP | K617T | 35011 |
| 8549 | A | G | 0.98 | VP30 | R14G | 30390 |
| 9690 | A | T | 0.97 | VP30 |  | 73597 |
| 9697 | A | C | 0.88 | VP30 |  | 68562 |
| 9698 | G | T | 0.87 | VP30 |  | 68992 |
| 9705 | A | T | 0.93 | VP30 |  | 63785 |
| 9824 | A | G | 0.54 |  |  | 67238 |
| 10833 | G | A | 0.57 | VP24 | R163K | 42279 |
| 10845 | T | A | 0.95 | VP24 | L167Q | 47557 |
| 11498 | G | A | 1.21 | VP24 |  | 43040 |
| 11695 | T | C | 1.41 | L | N38 | 41717 |
| 13001 | A | G | 0.59 | L | I480V | 43053 |
| 13234 | A | T | 0.50 | L | S551 | 39465 |
| 13240 | A | T | 0.59 | L | K553N | 36367 |
| 13497 | C | T | 0.57 | L | A639V | 69958 |
| 14043 | G | A | 1.41 | L | R821K | 47806 |
| 14412 | A | G | 1.78 | L | E944G | 49799 |
| 17507 | G | T | 0.66 | L | D1976Y | 46240 |
| 17510 | A | C | 0.58 | L | N1977H | 45945 |
| 18259 | T | G | 0.83 | L |  | 41881 |
| 18528 | T | C | 1.03 |  |  | 28861 |
| 18530 | A | T | 0.98 |  |  | 29397 |
| 18532 | G | A | 0.97 |  |  | 29743 |
| 18688 | A | G | 1.90 |  |  | 34663 |
| 18827 | G | C | 0.61 |  |  | 3908 |
| 18833 | G | T | 0.70 |  |  | 3573 |
| 18836 | A | C | 0.63 |  |  | 3331 |
| 18842 | G | C | 0.61 |  |  | 3133 |

We then characterized variation across passages in JK and HeLa cells. Taking into account all replicates and all passages, we detected a mean of 89 SNVs (σ = 31) for passages in HeLa cells and a mean of 51 SNVs (σ = 19) for passages in JK cells Table 1, Fig. 5A). Considering only nonsynonymous variants that were not already present in the inoculum, we detected a mean of 15 (σ = 15) for all replicates and all passages in HeLa cells and a mean of (σ = 7) for all replicates and all passages in JK cells (Table 1, Fig. 5B).

**FIG 5:**
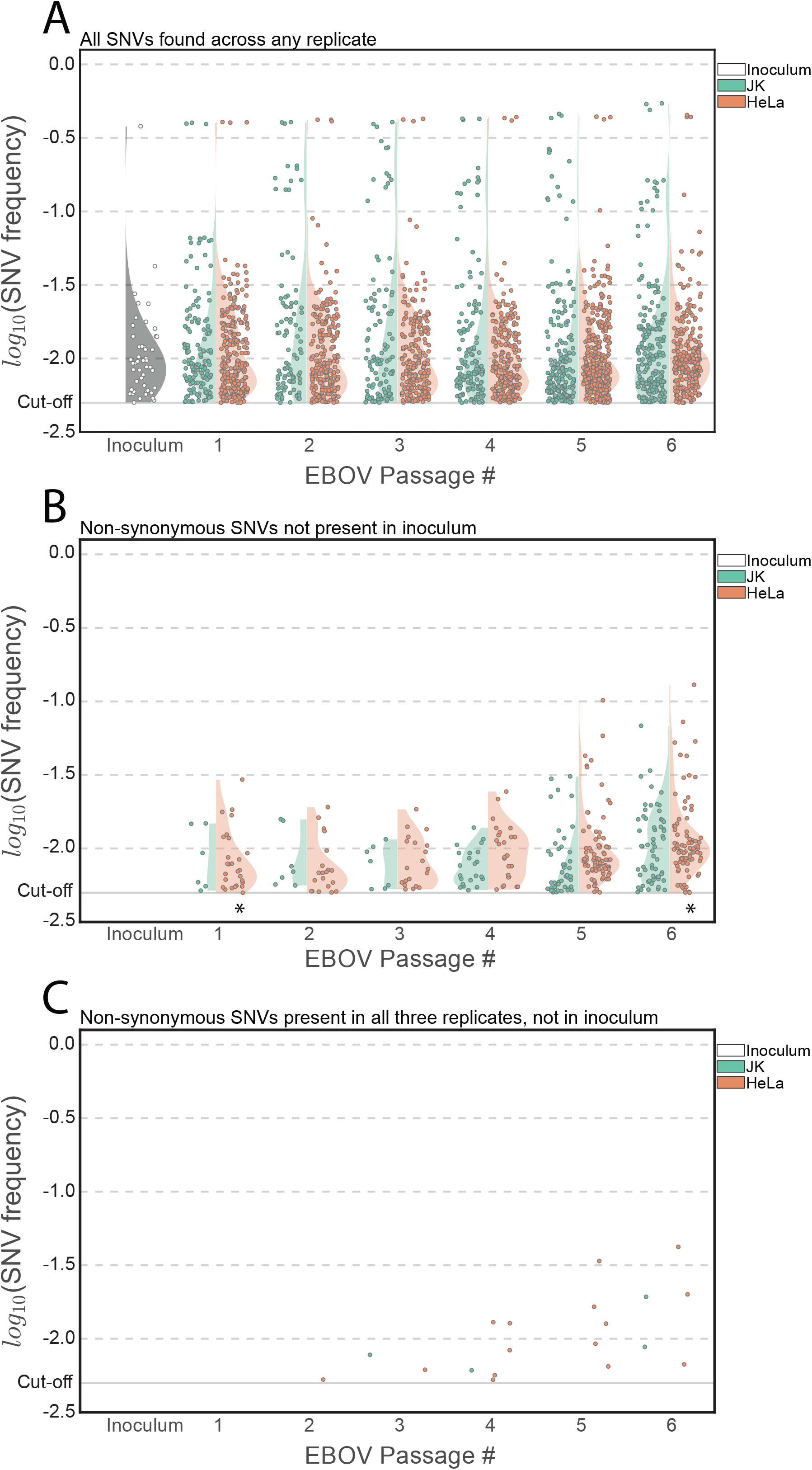
Graphs of EBOV passages vs. log10 (allele frequency) of single nucleotide variants (SNVs). Each SNV found in each passage was plotted as its log10 (allele frequency). A: Frequency of all SNVs from each replicate. B: Frequency of non-synonymous variants from each replicate that were not found in the inoculum. C: Non-synonymous variants found in all three replicates, but not the inoculum, were plotted as a single point with their mean frequency. Inoculum was a single replicate whereas all other passages were pooled triplicates, except for C. JK cells: green; HeLa cells: orange.

To determine whether there was a change in the distribution of allele frequencies associated with EBOV SNVs detected as a function of passage or host (boa constrictor vs. human) cell, we focused on a comparison of the first and last EBOV passages. The mean allele frequency associated with non-synonymous SNVs not found in the inoculum for EBOV grown in HeLa cells was 0.009 and 0.015 in in the first passage and passage 6, respectively. The difference between these passages was statistically significant (KS-test, p = 0.00051 < 0. Holm-Bonferroni adjusted p). However, the difference in distributions of allele frequencies associated with non-synonymous variants not found in the inoculum for EBOV grown in JK cells was not significant (KS-test, p = 0.41710 > 0.01 Holm-Bonferroni adjusted p).

We also compared the distribution of allele frequencies associated with nonsynonymous variants not found in the EBOV inoculum between the two host cells at the last passage. The difference between their means was relatively small (HeLa, JK mean = 0.015, 0.012, respectively), and the difference between these distributions was not statistically significant (KS-test, p = 0.0131 > 0.0083 Holm–Bonferroni adjusted p).

To further increase the stringency of our criteria for identifying biologically relevant EBOV variants, we considered only non-synonymous variants present in all three biological replicates for each passage from each host cell that were not present, or at a frequency below the limit of detection in the inoculum (Table 1, Fig. 5C). We detected a mean of 3 non-synonymous SNVs (σ = 2) across all passages in HeLa and a mean of 1 non-synonymous SNVs (σ = 1) across all passages in JK. We were unable to detect any EBOV SNVs that met these criteria for the first passage in either cell type. For JK passages, EBOV SNVs that met these criteria were only detected in passages 3, 4, and 6. In the case of passage 6 we did not find any statistical significance between the distributions of allele frequencies of SNVs found in the HeLa passage vs. the JK passage (KS-test p = 0.4249 vs. 0.05).

Finally, we implemented a rigorous simulation-based test for neutral evolution of EBOV that takes into account sequencing error, sampling error, and an estimated demographic model representing the passages in our experiments. We found numerous variants that deviate from neutral expectations (14,473 sites in JK and 15,028 sites in HeLa). However, as discussed above, nearly all of these variants experienced extremely small changes in allele frequency. To estimate the strength of selection operating on EBOV in each cell line, we implemented a deterministic fitness model and applied it to each site in turn. We found that the estimated selection coefficients are small (Fig. 4B, and Table S1).

Together, these data indicate that EBOV can replicate in boa constrictor cells for prolonged times/passages without requiring major genomic adaptations.

### Weak positive selection operates on the Ebola virus genome during passaging

To identify EBOV genomic sites undergoing positive selection in JK or HeLa cells, we first excluded sites with total read coverage that was not within two standard deviations of the genome-wide mean (calculated by first averaging the total reads across the three replicates for each passage and then averaging all passages). After filtering, a total of 17,924 sites and 17, sites, covering 95% of the genome, were retained for EBOV passaged on HeLa and JK cells respectively. Only three EBOV genomic sites had a change in allele frequency of at least 10%, all of which were identified in JK cell-grown virus (Figure 5C, Table S1): nucleotide positions 5,780 (located in VP40 5’ UTR), 7,669 (preGP T544I), and 18,016 (L, sysnonymous mutation). In HeLa cells, all allele frequency changes were less than 7% (Table S1). Using a deterministic model of positive selection (see supplemental methods), we estimate that the selection coefficient at all sites in the EBOV genome (across both HeLa and JK cells) was less than 12%. These data suggest that weak selection can be identified in the EBOV genome over passages (particularly in JK cells; see supplemental methods for statistical test results), but that very little adaptation is necessary to successfully passage EBOV in either cell type.

### Passage of Ebola virus in either Boa constrictor cells or HeLa does not lead to appreciable production of defective interfering genomes

The presence of DI particles has been noted with EBOV in grivet (*Chlorocebus aethiops*) kidney epithelial Vero E6 cell culture, but they remain poorly understood with only a single paper published on EBOV DI genome characterization (37). Viral DI particles often contain genomes with long deletions or genomic re-arrangements that presumably arise through errors in replication by, for instance, template switching (38). To detect the presence of EBOV genomic sequences with deletions that would likely yield DI particles, we quantified the insertion distance between sequence pairs for EBOV infecting both JK and HeLa cells across all passages and replicates. The EBOV inoculum featured 0.0276% of reads that were consistent with internal genomic deletions. We detected a low level of putative deletion sequences in both cell types (0.0780% σ= 0.0535 for HeLa cells and 0.0234% σ= 0.00480 for JK cells) in all passages and replicates distributed across the EBOV genome (Table 1, Fig. S1). By the final passage, this value changed to 0.0552% σ= 0.00340 and 0.0206% σ= 0.00185 for HeLa and JK cells, respectively. In this analysis, we can not rule out the possibility of internal deletions produced during sequencing library preparation, and thus these measurements are likely to be over estimates. Regardless, this analysis indicates that sequences consistent with the presence of DI particles could be detected, but only at very low frequencies.

## DISCUSSION

The natural reservoir of EBOV and all other ebolaviruses remains unclear. Marburgviruses have been isolated from wild Ugandan Egyptian rousettes (*Rousettus aegyptiacus*) and also have been used to infect these bats experimentally (11, 15, 16). Such findings have not been reported for ebolaviruses, thereby raising the possibility that marburgviruses and ebolaviruses may differ in host tropism and may even infect animals of different orders (11-14, 39). Experimental filovirus inoculations into taxonomically diverse animals to determine host tropism have only rarely been reported. These experiments suggest that all isolated filoviruses can infect and are frequently lethal for various nonhuman primates (common marmosets [*Callithrix jacchus*], common squirrel monkeys [*Saimiri sciureus*], crab-eating macaques [*Macaca fascicularis*], grivets [*Chlorocebus aethiops*], hamadryas baboons [*Papio hamadryas*], rhesus monkeys [*Macaca mulatta*]) and domestic ferrets (*Mustela putorius furo*); and that most filoviruses can be adapted in the laboratory to infect and kill various rodents (golden hamsters [*Mesocricetus auratus*], guinea pigs [*Cavia porcellus*], laboratory mice); and that some filoviruses can infect domestic pigs (*Sus scrofa*). Various plants, goats (*Capra hircus*), horses (*Equus caballus*), and red sheep (*Ovis aries*) were found to be resistant to experimental filovirus infection (summarized in (3, 40, 41)).

In 2001, a possible genetic link between mammalian arenaviruses (family *Arenaviridae*, genus *Mammarenavirus*) and the mononegaviral filoviruses was suggested based on similarities between arenaviral and filoviral glycoproteins (17). This possible link was further substantiated by the structural characterization of the glycoprotein from a newly discovered snake reptarenavirus (genus *Reptarenavirus*) (42). Filoviral glycoproteins engage endosomal mammalian Niemann-Pick disease, type C1 protein (NPC1) to gain entry into host cell (12, 43). Interestingly, Russell’s viper (*Daboia russellii*) cells do not support EBOV entry and Russell’s viper NPC1 does not bind to the EBOV glycoprotein. This deficiency was traced to a single amino acid (Y503) that, when changed to the analogous human residue (Y503F), causes Russel’s viper cells to become susceptible to EBOV infection (44). Although the boa constrictor genome has been assembled, it has not been annotated (REF PMID 23870653). We used a comparative alignment approach and mapping of transcriptome-derived short sequence reads to predict the boa constrictor NPC1 protein sequence (Genbank XXXXXXX). The predicted boa constrictor NPC1 has an Phe residue at the critical position (F517, homologous to F503 in human NPC1), which is consistent with boa constrictor cell susceptibility to EBOV infection. This supports the possibility that NPC1 from snakes of certain species may have been subject to selection by viruses with filovirus-like glycoproteins (44).

We aimed to further explore the potential genetic link between filoviruses and reptarenaviruses. Reptarenaviruses are known to infect captive boid snakes (pythons and boas) (18, 45-47). We tested whether boa constrictor JK cells are naturally capable of supporting EBOV or MARV infection. While MARV infection was unsuccessful, JK cells supported EBOV replication over six passages in the absence of major genomic adaptation. Only one genomic position, 7669, (EBOV preGP T544I) switched major alleles (38% to 52%). After maturation of the glycoprotein precursor, this residue resides in the preGP cleavage product GP2. The residue is a critical structural determinant of the EBOV GP2 fusion loop, which mediates fusion of the filovirion membrane with the host-cell membrane to initiate virion entry (48). Both alleles, Thr and Ile, have been identified in different EBOV isolates. For instance, isolates of the EBOV Makona variant (49), which caused the 2013–2016 Western African EVD outbreak, almost exclusively encode Thr at preGP position 544 (4-10), whereas a 1976 EBOV Yambuku variant isolate encodes the Ile allele (50). Curiously, Ile is also encoded at the homologous position in the genome of RESTV (51, 52), which has not yet been associated with human infections. We detected weak positive selection favoring the Ile allele in the EBOV passages in JK cells, suggesting this allele provides a fitness advantage over Thr for infection in JK cells, but the mechanistic reason for this selection remains to be determined. In contrast to the successful infection of JK cells with EBOV, these cells were unable to support productive MARV infection. Uncovering the molecular underpinnings of this difference could increase our understanding of filovirus tropism. Importantly, EBOV infection of JK cells occurred in the absence of CPE, an observation that also has only rarely been reported (53). These observations raise the possibility that ebolaviruses and marburgviruses could sub-clinically and/or persistently infect disparate hosts, possibly even of different animal orders (e.g., mammals vs. reptiles). Additional non-mammalian cell lines should therefore be screened for filovirus susceptibility to aid the search for natural filovirus hosts, possibly followed by experimental animal inoculations or exposures. These experiments further suggest that additional genetic dissection of MARV will reveal the underlying determinants that prevent JK cell infection.

## ACKNOWLEDGMENTS

We thank Laura Bollinger (NIH/NIAID Integrated Research Facility at Fort Detrick, Frederick, MD, USA) for critically editing the manuscript.

## FUNDING INFORMATION

This work was supported by the Howard Hughes Medical Institute, and in part through Battelle Memorial Institute’s prime contract with the US National Institute of Allergy and Infectious Diseases (NIAID) under Contract No. HHSN272200700016I, and by the US National Human Genome Research Institute (R01 HG007644) to RDH. A subcontractor to Battelle Memorial Institute who performed this work is: J.H.K., an employee of Tunnell Government Services, Inc.

The views and conclusions contained in this document are those of the authors and should not be interpreted as necessarily representing the official policies, either expressed or implied, of the US Department of the Army, the US Department of Defense, the US Department of Health and Human Services, or of the institutions and companies affiliated with the authors.

## SUPPLEMENTAL DATA

**Supplemental figure S1**: Mapping location of sequencing read pairs of possible defective interfering genomes. For each properly paired end sequencing reads, the starting position of read 1 mapped onto the EBOV reference genome is plotted against the starting position of read 2 mapped onto the EBOV reference genome. The left window is 1 × 106 read pairs randomly sampled, without replacement, from the inoculum. The center window is 1 × 106 read pairs randomly sampled, without replacement, from the all the passages in HeLa cells. The right window is 1 × 106 read pairs randomly sampled, without replacement, from the all the passages in JK cells. Each read pair (black dot) has an alpha value (opacity) of 0.1.

**Supplemental figure S2**: Population sizes over time. Depicts the estimated and observed population sizes over time. The continuous red lines show the population size that was estimated from the observed population sizes (black points) at six equally spaced time points. Significant bottlenecks occurred with each passage. Estimates were determined separately for each cell type and trial.

**Supplemental table S1**: Estimated selection coefficients of significant positively selected sites across the EBOV genome. Each row represents a genome position with their allele, the respective average (across the 3 replicates) selection coefficient, average (across the 3 replicates) change in allele frequency between starting and ending passages, and the p value. p values were calculated using Fisher’s method.

**Supplemental table S2:** MARV passage quantification. Each element in the table represents MARV genomes per ml in the supernatant found by qRT-PCR of each sample. Samples that did not cross threshold after 40 cycles are labeled as not detected, ND.

## REFERENCES

1. Kuhn JH. 2015. Ebolavirus and marburgvirus Infections, p 1323–1329. In Kasper DL, Fauci AS, Hauser SL, Longo DL, Jameson JL, Loscalzo J (ed), Harrison’s Principles of Internal Medicine, 19th ed, vol 2. McGraw-Hill Education, Columbus, Ohio, USA.

2. World Health Organization. 2016. Ebola situation reports. http://apps.who.int/ebola/ebola-situation-reports.

3. Kuhn JH. 2008. Filoviruses. A compendium of 40 years of epidemiological, clinical, and laboratory studies. Archives of Virology Supplementum, vol. 20. Springer Wien New York, Vienna, Austria.

4. Park DJ, Dudas G, Wohl S, Goba A, Whitmer SLM, Andersen KG, Sealfon RS, Ladner JT, Kugelman JR, Matranga CB, Winnicki SM, Qu J, Gire SK, Gladden-Young A, Jalloh S, Nosamiefan D, Yozwiak NL, Moses LM, Jiang P-P, Lin AE, Schaffner SF, Bird B, Towner J, Mamoh M, Gbakie M, Kanneh L, Kargbo D, Massally JLB, Kamara FK, Konuwa E, Sellu J, Jalloh AA, Mustapha I, Foday M, Yillah M, Erickson BR, Sealy T, Blau D, Paddock C, Brault A, Amman B, Basile J, Bearden S, Belser J, Bergeron E, Campbell S, Chakrabarti A, Dodd K, Flint M, Gibbons A, et al. 2015. Ebola virus epidemiology, transmission, and evolution during seven months in Sierra Leone. Cell 161:1516–1526.

5. Ladner JT, Wiley MR, Mate S, Dudas G, Prieto K, Lovett S, Nagle ER, Beitzel B, Gilbert ML, Fakoli L, Diclaro JW 2nd, Schoepp RJ, Fair J, Kuhn JH, Hensley LE, Park DJ, Sabeti PC, Rambaut A, Sanchez-Lockhart M, Bolay FK, Kugelman JR, Palacios G. 2015. Evolution and spread of Ebola Virus in Liberia, 2014-2015. Cell Host Microbe 18:659–669.

6. Gire SK, Goba A, Andersen KG, Sealfon RSG, Park DJ, Kanneh L, Jalloh S, Momoh M, Fullah M, Dudas G, Wohl S, Moses LM, Yozwiak NL, Winnicki S, Matranga CB, Malboeuf CM, Qu J, Gladden AD, Schaffner SF, Yang X, Jiang P-P, Nekoui M, Colubri A, Coomber MR, Fonnie M, Moigboi A, Gbakie M, Kamara FK, Tucker V, Konuwa E, Saffa S, Sellu J, Jalloh AA, Kovoma A, Koninga J, Mustapha I, Kargbo K, Foday M, Yillah M, Kanneh F, Robert W, Massally JLB, Chapman SB, Bochicchio J, Murphy C, Nusbaum C, Young S, Birren BW, Grant DS, Scheiffelin JS, et al. 2014. Genomic surveillance elucidates Ebola virus origin and transmission during the 2014 outbreak. Science 345:1369–1372.

7. Carroll MW, Matthews DA, Hiscox JA, Elmore MJ, Pollakis G, Rambaut A, Hewson R, García-Dorival I, Bore JA, Koundouno R, Abdellati S, Afrough B, Aiyepada J, Akhilomen P, Asogun D, Atkinson B, Badusche M, Bah A, Bate S, Baumann J, Becker D, Becker-Ziaja B, Bocquin A, Borremans B, Bosworth A, Boettcher JP, Cannas A, Carletti F, Castilletti C, Clark S, Colavita F, Diederich S, Donatus A, Duraffour S, Ehichioya D, Ellerbrok H, Fernandez-Garcia MD, Fizet A, Fleischmann E, Gryseels S, Hermelink A, Hinzmann J, Hopf-Guevara U, Ighodalo Y, Jameson L, Kelterbaum A, Kis Z, Kloth S, Kohl C, Korva M, et al. 2015. Temporal and spatial analysis of the 2014-2015 Ebola virus outbreak in West Africa. Nature 524:97–101.

8. Simon-Loriere E, Faye O, Faye O, Koivogui L, Magassouba N, Keita S, Thiberge J-M, Diancourt L, Bouchier C, Vandenbogaert M, Caro V, Fall G, Buchmann JP, Matranga CB, Sabeti PC, Manuguerra J-C, Holmes EC, Sall AA. 2015. Distinct lineages of Ebola virus in Guinea during the 2014 West African epidemic. Nature 524:102–104.

9. Tong Y-G, Shi W-F, Liu D, Qian J, Liang L, Bo X-C, Liu J, Ren H-G, Fan H, Ni M, Sun Y, Jin Y, Teng Y, Li Z, Kargbo D, Dafae F, Kanu A, Chen C-C, Lan Z-H, Jiang H, Luo Y, Lu H-J, Zhang X-G, Yang F, Hu Y, Cao Y-X, Deng Y-Q, Su H-X, Sun Y, Liu W-S, Wang Z, Wang C-Y, Bu Z-Y, Guo Z-D, Zhang L-B, Nie W-M, Bai C-Q, Sun C-H, An X-P, Xu P-S, Zhang X-L-L, Huang Y, Mi Z-Q, Yu D, Yao H-W, Feng Y, Xia Z-P, Zheng X-X, Yang S-T, Lu B, et al. 2015. Genetic diversity and evolutionary dynamics of Ebola virus in Sierra Leone. Nature 524:93–96.

10. Baize S, Pannetier D, Oestereich L, Rieger T, Koivogui L, Magassouba NF, Soropogui B, Sow MS, Keïta S, De Clerck H, Tiffany A, Dominguez G, Loua M, Traoré A, Kolié M, Malano ER, Heleze E, Bocquin A, Mély S, Raoul H, Caro V, Cadar D, Gabriel M, Pahlmann M, Tappe D, Schmidt-Chanasit J, Impouma B, Diallo AK, Formenty P, Van Herp M, Günther S. 2014. Emergence of Zaire Ebola virus disease in Guinea. N Engl J Med 371:1418–1425.

11. Jones MEB, Schuh AJ, Amman BR, Sealy TK, Zaki SR, Nichol ST, Towner JS. 2015. Experimental inoculation of Egyptian rousette bats (Rousettus aegyptiacus) with viruses of the Ebolavirus and Marburgvirus genera. Viruses 7:3420–3442.

12. Wahl-Jensen V, Radoshitzky SR, de Kok-Mercado F, Taylor SL, Bavari S, Jahrling PB, Kuhn JH. 2013. Role of rodents and bats in human viral hemorrhagic fevers, p 99– 127. In Singh SK, Ruzek D (ed), Viral Hemorrhagic Fevers doi:10.1201/b15172-9. Taylor & Francis/CRC Press, Boca Raton, Florida, USA.

13. Paweska JT, Storm N, Grobbelaar AA, Markotter W, Kemp A, Jansen van Vuren P. 2016. Experimental inoculation of Egyptian fruit bats (Rousettus aegyptiacus) with Ebola virus. Viruses 8:29.

14. Leendertz SAJ, Gogarten JF, Düx A, Calvignac-Spencer S, Leendertz FH. 2016. Assessing the evidence supporting fruit bats as the primary reservoirs for Ebola viruses. Ecohealth 13:18–25.

15. Towner JS, Amman BR, Sealy TK, Carroll SA, Comer JA, Kemp A, Swanepoel R, Paddock CD, Balinandi S, Khristova ML, Formenty PBH, Albarino CG, Miller DM, Reed ZD, Kayiwa JT, Mills JN, Cannon DL, Greer PW, Byaruhanga E, Farnon EC, Atimnedi P, Okware S, Katongole-Mbidde E, Downing R, Tappero JW, Zaki SR, Ksiazek TG, Nichol ST, Rollin PE. 2009. Isolation of genetically diverse Marburg viruses from Egyptian fruit bats. PLoS Pathog 5:e1000536.

16. Amman BR, Carroll SA, Reed ZD, Sealy TK, Balinandi S, Swanepoel R, Kemp A, Erickson BR, Comer JA, Campbell S, Cannon DL, Khristova ML, Atimnedi P, Paddock CD, Kent Crockett RJ, Flietstra TD, Warfield KL, Unfer R, Katongole-Mbidde E, Downing R, Tappero JW, Zaki SR, Rollin PE, Ksiazek TG, Nichol ST, Towner JS. 2012. Seasonal pulses of Marburg virus circulation in juvenile Rousettus aegyptiacus bats coincide with periods of increased risk of human infection. PLoS Pathog 8:e1002877.

17. Gallaher WR, DiSimone C, Buchmeier MJ. 2001. The viral transmembrane superfamily: possible divergence of arenavirus and filovirus glycoproteins from a common RNA virus ancestor. BMC Microbiol 1:1.

18. Stenglein MD, Sanders C, Kistler AL, Ruby JG, Franco JY, Reavill DR, Dunker F, Derisi JL. 2012. Identification, characterization, and in vitro culture of highly divergent arenaviruses from boa constrictors and annulated tree boas: candidate etiological agents for snake inclusion body disease. MBio 3:e00180–00112.

19. Kugelman JR, Rossi CA, Wiley MR, Ladner JT, Nagle ER, Pfeffer BP, Garcia K, Prieto K, Wada J, Kuhn JH, Palacios G. 2016. Informing the historical record of experimental nonhuman primate infections with Ebola virus: genomic characterization of USAMRIID Ebola virus/H.sapiens-tc/COD/1995/Kikwit-9510621 challenge stock “R4368” and its replacement “R4415”. PLoS One 11:e0150919.

20. Smith DH, Johnson BK, Isaacson M, Swanapoel R, Johnson KM, Killey M, Bagshawe A, Siongok T, Keruga WK. 1982. Marburg-virus disease in Kenya. Lancet 1:816–820.

21. Shurtleff AC, Biggins JE, Keeney AE, Zumbrun EE, Bloomfield HA, Kuehne A, Audet JL, Alfson KJ, Griffiths A, Olinger GG, Bavari S. 2012. Standardization of the filovirus plaque assay for use in preclinical studies. Viruses 4:3511–3530.

22. Shurtleff AC, Bloomfield HA, Mort S, Orr SA, Audet B, Whitaker T, Richards MJ, Bavari S. 2016. Validation of the filovirus plaque assay for use in preclinical studies. Viruses 8:113.

23. Moe JB, Lambert RD, Lupton HW. 1981. Plaque assay for Ebola virus. J Clin Microbiol 13:791–793.

24. Radoshitzky SR, Dong L, Chi X, Clester JC, Retterer C, Spurgers K, Kuhn JH, Sandwick S, Ruthel G, Kota K, Boltz D, Warren T, Kranzusch PJ, Whelan SPJ, Bavari S. 2010. Infectious Lassa virus, but not filoviruses, is restricted by BST-5422/tetherin. J Virol 84:10569–10580.

25. Hindson BJ, Ness KD, Masquelier DA, Belgrader P, Heredia NJ, Makarewicz AJ, Bright IJ, Lucero MY, Hiddessen AL, Legler TC, Kitano TK, Hodel MR, Petersen JF, Wyatt PW, Steenblock ER, Shah PH, Bousse LJ, Troup CB, Mellen JC, Wittmann DK, Erndt NG, Cauley TH, Koehler RT, So AP, Dube S, Rose KA, Montesclaros L, Wang S, Stumbo DP, Hodges SP, Romine S, Milanovich FP, White HE, Regan JF, Karlin-Neumann GA, Hindson CM, Saxonov S, Colston BW. 2011. High-throughput droplet digital PCR system for absolute quantitation of DNA copy number. Anal Chem 83:8604–8610.

26. Stenglein MD, Jacobson ER, Wozniak EJ, Wellehan JF, Kincaid A, Gordon M, Porter BF, Baumgartner W, Stahl S, Kelley K, Towner JS, DeRisi JL. 2014. Ball python nidovirus: a candidate etiologic agent for severe respiratory disease in Python regius. MBio 5:e01484–01414.

27. Wilson MR, Fedewa G, Stenglein MD, Olejnik J, Rennick LJ, Nambulli S, Feldmann F, Duprex WP, Connor JH, Mühlberger E, DeRisi JL, Chia N. 2016. Multiplexed metagenomic deep sequencing to analyze the composition of high-priority pathogen reagents. mSystems 1:e00058–00016.

28. Ruby JG, Bellare P, Derisi JL. 2013. PRICE: software for the targeted assembly of components of (Meta) genomic sequence data. G3 (Bethesda) 3:865–880.

29. Wu TD, Nacu S. 2010. Fast and SNP-tolerant detection of complex variants and splicing in short reads. Bioinformatics 26:873–881.

30. Wilm A, Aw PPK, Bertrand D, Yeo GHT, Ong SH, Wong CH, Khor CC, Petric R, Hibberd ML, Nagarajan N. 2012. LoFreq: a sequence-quality aware, ultra-sensitive variant caller for uncovering cell-population heterogeneity from high-throughput sequencing datasets. Nucleic Acids Res 40:11189–11201.

31. Pérez F, Granger BE. 2007. IPython: a system for interactive scientific computing. Computing in Science & Engineering 9:21–29.

32. McKinney W. 2011. pandas: a Foundational Python Library for Data Analysis and Statistics.

33. Hunter JD. 2007. Matplotlib: a 2D graphics environment. Computing in Science & Engineering 9:90–95.

34. Waskom M, Botvinnik O, drewokane, Hobson P, Halchenko Y, Lukauskas S, Warmenhoven J, Cole JB, Hoyer S, Vanderplas J, gkunter, Villalba S, Quintero E, Martin M, Miles A, Meyer K, Augspurger T, Yarkoni T, Bachant P, Evans C, Fitzgerald C, Nagy T, Ziegler E, Megies T, Wehner D, St-Jean S, Coelho LP, Hitz G, Lee A, Rocher L. 2016. seaborn: v0.7.0 (January 2016). zenodo doi:10.5281/zenodo.45133.

35. Groseth A, Marzi A, Hoenen T, Herwig A, Gardner D, Becker S, Ebihara H, Feldmann H. 2012. The Ebola virus glycoprotein contributes to but is not sufficient for virulence in vivo. PLoS Pathog 8:e1002847.

36. Shabman RS, Jabado OJ, Mire CE, Stockwell TB, Edwards M, Mahajan M, Geisbert TW, Basler CF. 2014. Deep sequencing identifies noncanonical editing of Ebola and Marburg virus RNAs in infected cells. MBio 5:e02011.

37. Calain P, Monroe MC, Nichol ST. 1999. Ebola virus defective interfering particles and persistent infection. Virology 262:114–128.

38. Lazzarini RA, Keene JD, Schubert M. 1981. The origins of defective interfering particles of the negative-strand RNA viruses. Cell 26:145–154.

39. Jensen Leendertz SA. 2016. Testing new hypotheses regarding ebolavirus reservoirs. Viruses-Basel 8:30.

40. Burk R, Bollinger L, Johnson JC, Wada J, Radoshitzky SR, Palacios G, Bavari S, Jahrling PB, Kuhn JH. 2016. Neglected filoviruses. FEMS Microbiol Rev 40:494–519.

41. Swanepoel R, Leman PA, Burt FJ, Zachariades NA, Braack LEO, Ksiazek TG, Rollin PE, Zaki SR, Peters CJ. 1996. Experimental inoculation of plants and animals with Ebola virus. Emerg Infect Dis 2:321–325.

42. Koellhoffer JF, Dai Z, Malashkevich VN, Stenglein MD, Liu Y, Toro R J SH, Chandran K, DeRisi JL, Almo SC, Lai JR. 2014. Structural characterization of the glycoprotein GP2 core domain from the CAS virus, a novel arenavirus-like species. J Mol Biol 426:1452–1468.

43. Côte M, Misasi J, Ren T, Bruchez A, Lee K, Filone CM, Hensley L, Li Q, Ory D, Chandran K, Cunningham J. 2011. Small molecule inhibitors reveal Niemann-Pick C is essential for Ebola virus infection. Nature 477:344–348.

44. Ndungo E, Herbert AS, Raaben M, Obernosterer G, Biswas R, Miller EH, Wirchnianski AS, Carette JE, Brummelkamp TR, Whelan SP, Dye JM, Chandran K. 2016. A single residue in Ebola virus receptor NPC1 influences cellular host range in reptiles. mSphere 1:e00007–00016.

45. Bodewes R, Kik MJL, Raj VS, Schapendonk CME, Haagmans BL, Smits SL, Osterhaus ADME. 2013. Detection of novel divergent arenaviruses in boid snakes with inclusion body disease in The Netherlands. J Gen Virol 94:1206–1210.

46. Hepojoki J, Salmenperä P, Sironen T, Hetzel U, Korzyukov Y, Kipar A, Vapalahti O. 2015. Arenavirus coinfections are common in snakes with boid inclusion body disease. J Virol 89:8657–8660.

47. Stenglein MD, Jacobson ER, Chang L-W, Sanders C, Hawkins MG, Guzman DS-M, Drazenovich T, Dunker F, Kamaka EK, Fisher D, Reavill DR, Meola LF, Levens G, DeRisi JL. 2015. Widespread recombination, reassortment, and transmission of unbalanced compound viral genotypes in natural arenavirus infections. PLoS Pathog 11:e1004900.

48. Gregory SM, Larsson P, Nelson EA, Kasson PM, White JM, Tamm LK. 2014. Ebolavirus entry requires a compact hydrophobic fist at the tip of the fusion loop. J Virol 88:6636–6649.

49. Kuhn JH, Andersen KG, Baize S, Bào Y, Bavari S, Berthet N, Blinkova O, Brister JR, Clawson AN, Fair J, Gabriel M, Garry RF, Gire SK, Goba A, Gonzalez J-P, Günther S, Happi CT, Jahrling PB, Kapetshi J, Kobinger G, Kugelman JR, Leroy EM, Maganga GD, Mbala PK, Moses LM, Muyembe-Tamfum J-J, Magassouba NF, Nichol ST, Omilabu SA, Palacios G, Park DJ, Paweska JT, Radoshitzky SR, Rossi CA, Sabeti PC, Schieffelin JS, Schoepp RJ, Sealfon R, Swanepoel R, Towner JS, Wada J, Wauquier N, Yozwiak NL, Formenty P. 2014. Nomenclature- and database-628compatible names for the two Ebola virus variants that emerged in Guinea and the Democratic Republic of the Congo in 2014. Viruses 6:4760–4799.

50. Kuhn JH, Lofts LL, Kugelman JR, Smither SJ, Lever MS, van der Groen G, Johnson KM, Radoshitzky SR, Bavari S, Jahrling PB, Towner JS, Nichol ST, Palacios G. 2014. Reidentification of Ebola virus E718 and ME as Ebola virus/H.sapiens-tc/COD/1976/Yambuku-Ecran. Genome Announc 2.

51. Groseth A, Ströher U, Theriault S, Feldmann H. 2002. Molecular characterization of an isolate from the 1989/90 epizootic of Ebola virus Reston among macaques imported into the United States. Virus Res 87:155–163.

52. Ikegami T, Calaor AB, Miranda ME, Niikura M, Saijo M, Kurane I, Yoshikawa Y, Morikawa S. 2001. Genome structure of Ebola virus subtype Reston: differences among Ebola subtypes. Arch Virol 146:2021–2027.

53. van der Groen G, Webb P, Johnson K, Lange JV, Lindsay H, Eliott L. 1978. Growth of Lassa and Ebola viruses in different cell lines, p 255–260. In Pattyn SR (ed), Ebola virus haemorrhagic fever. Elsevier/North-Holland Biomedical Press, Amsterdam, The Netherlands.

